# Protease-Resistant Azapeptide GLP-1 Analogue Improves Metabolic Control in Diet-Induced Obesity

**DOI:** 10.1101/2025.05.09.653092

**Authors:** Mingzhu He, Kai Fan Cheng, Sonya VanPatten, Marcelo D. T. Torres, Bayan Al Jabari, Ibrahim T. Mughrabi, Borja Ballarín-González, Myoungsun Son, Cesar de la Fuente-Nunez, Yousef Al-Abed

**Affiliations:** The Feinstein Institutes for Medical Research, Northwell Health, Manhasset, NY, USA; Machine Biology Group, Departments of Psychiatry and Microbiology, Institute for Biomedical Informatics, Institute for Translational Medicine and Therapeutics, Perelman School of Medicine, University of Pennsylvania, Philadelphia, PA, USA; Departments of Bioengineering and Chemical and Biomolecular Engineering, School of Engineering and Applied Science, University of Pennsylvania, Philadelphia, PA, USA; Department of Chemistry, School of Arts and Sciences, University of Pennsylvania, Philadelphia, PA, USA; Penn Institute for Computational Science, University of Pennsylvania, Philadelphia, PA, USA; Institute of Bioelectronic Medicine, Feinstein Institutes for Medical Research, Northwell Health, Manhasset, NY USA; Novo Nordisk A/S, Måløv, Denmark; Institute of Molecular Medicine, Feinstein Institutes for Medical Research, Northwell Health, Manhasset, NY, USA; Donald and Barbara Zucker School of Medicine at Hofstra/Northwell, Northwell Health, Hempstead, NY, USA

**Keywords:** azapeptide, GLP-1 receptor agonist (GLP-1RA), high-fat diet-induced obesity (HF-DIO)

## Abstract

Peptide therapeutics are limited by rapid proteolysis and short half-lives. Azapeptides, created by replacing one or more α-carbon(s) on the peptide backbone with nitrogen atom(s), offer a strategy to improve peptide stability while preserving functional efficiency, yet their clinical potential has remained underexplored. Herein, we report the design, synthesis, *in vitro* and *in vivo* evaluations of azapeptide-based glucagon-like peptide-1 receptor agonists (GLP-1RAs). Using a solid-phase synthesis platform, we generated GLP-1 analogues with aza-substitutions at protease-sensitive residues. The lead analogue, AzaA^8^/R^34^-GLP-1(AzaA8), resisted dipeptidyl peptidase-4 degradation (>24 h), maintained picomolar potency at the GLP-1 receptor (GLP-1R) signaling, and exhibited an extended plasma half-life in mice relative to unmodified controls. In lean mice, AzaA8 improved oral glucose tolerance, and in high-fat diet-induced obese mice, chronic administration reduced body weight, decreased leptin and insulin levels, and enhanced glucose handling without detectable inflammatory adverse effects. These findings demonstrate that a targeted aza-substitution yields a protease-stable, biologically active GLP-1RA with metabolic benefits, establishing azapeptides as a promising scaffold for next-generation incretin-based therapies in diabetes and obesity.

## Introduction

Obesity and type 2 diabetes are escalating global health crises that drive cardiovascular disease, fatty liver, and other metabolic complications^1^. Current therapies often show limited long-term efficacy, side effects, or require frequent administration, underscoring the need for safer, longer-acting interventions^2–4^. One successful class of treatments is the incretin-based peptide therapeutics, exemplified by glucagon-like peptide-1 receptor agonists (GLP-1RAs), which enhance insulin secretion and suppress appetite^5^.

GLP-1RAs such as semaglutide and liraglutide (**Fig. 1a, 1d**) have transformed the management of diabetes and obesity by improving glycemic control and inducing weight loss. However, native GLP-1 is rapidly inactivated by dipeptidyl peptidase-4 (DPP-4; **Fig. 1b**), and even clinically used analogues require extensive chemical modifications to modestly extend half-life^6^. For example, Ozempic® (semaglutide) retains the GLP-1(7–37) scaffold but incorporates an Ala8→α-aminoisobutyric acid (Aib) substitution to block DPP-4 cleavage, a Lys34→Arg substitution, and an 18-carbon fatty diacid attached at Lys26 to confer albumin binding and slow clearance. While these modifications are effective, they add complexity and highlight the need for alternative strategies to engineer peptides with intrinsic stability.

**Figure 1.** GLP-1/semaglutide engagement with GLP-1R and sequence schematics. **a)** Structural overlay of the GLP-1: GLP-1R complex (ligand, red; receptor, pink, PDBid: 6X18) and the semaglutide: GLP-1R complex (ligand, blue; receptor, light blue, PDBid: 7KI0), aligned on the receptor. The boxed inset enlarges the peptide–receptor interface to illustrate representative hydrogen bonds, salt bridges, and hydrophobic contacts. **b)** Linear map of R34-GLP-1 showing only residues that contact GLP-1R (highlighted and numbered); non-contact positions are gray. Schematic of glucagon-like peptide-1 (GLP-1) and analogues. Sequence and descriptive (abbreviated names) information for **c)** GLP-1 and R34-GLP-1, **d)** unlipidated semaglutide, semaglutide, and liraglutide, **e)** and azaGLP-1-based analogues. Red dashed lines represent the DPP-4 cleavage site. The protein and peptide structures depicted in panels **a** and **b** were created with PyMOL Molecular Graphics System, v.3.0 (Schrödinger).

Azapeptides, in which α-carbon(s) is replaced by nitrogen(s) in the peptide backbone, offer one such strategy. This conservative substitution preserves the overall backbone geometry while rendering the modified bond resistant to protease cleavage^7,8^. Despite their potential to increase stability and half-life, azapeptides have seen limited clinical translation [goserelin (Zoladex®) is the sole FDA-approved example], in part due to historical synthetic challenges^9^. We recently developed a platform method that enables efficient incorporation of pre-activated aza-amino acid building blocks into peptides using standard solid-phase synthesis^10–13^, opening the door to explore azapeptide therapeutics more broadly.

Here, we applied this approach to a therapeutically relevant peptide hormone, designing GLP-1 analogues with aza-substitutions at known protease-sensitive positions. We identified a lead candidate, azaA8 (Ala8→aza-Ala in an Arg34-GLP-1 background), that resists DPP-4 degradation, retains low-picomolar receptor agonist potency, and produces significant metabolic benefits in both lean and diet-induced obese mice. These findings demonstrate that aza-substitution is a generalizable strategy for generating stable, efficacious incretin mimetics with translational potential for the treatment of diabetes and obesity.

## Results

### Azapeptide design and DPP-4 stability

To evaluate whether backbone aza-substitution(s) can stabilize a clinically relevant peptide hormone, we synthesized GLP-1(7–37) analogues bearing N-terminal aza-residues using our solid-phase platform^13^. The parent sequence was modified with a Lys34→Arg substitution (R34-GLP-1, as in semaglutide). We generated analogues with aza-substitutions at His7, Ala8, both His7 and Ala8 (with Ala8 optionally replaced by Aib in one variant), and at Glu9 (**Fig. 1c–e**), and benchmarked against unmodified R34-GLP-1, “unlipidated semaglutide” (Aib8), marketed GLP-1RAs (semaglutide, liraglutide, and exendin-4). All peptides were synthesized and purified using our standard solid-phase methods with aza-amino acid building blocks, and their identities and purities were confirmed by mass spectrometry (**Supplementary Table 1**)^13^.

In recombinant DPP-4 digestion assays, R34-GLP-1(7–37) was degraded within minutes (half-life, t_½_ ≈ 0.3 h). The protective Aib8 substitution in the unlipidated semaglutide control extended its half-life to ∼5.4 h (**Table 1** and **Supplementary Table 2**), whereas, all aza-modified analogues remained intact beyond 24 h, with no measurable loss after 48 h. Thus, replacement of His7, Ala8, or adjacent residues with aza-amino acids completely abolished cleavage at the canonical DPP-4 site, conferring a level of protection surpassing current clinical modifications.

**Table 1.** Composition, stability, pharmacokinetics, and receptor potency of GLP-1 analogues. Recombinant DPP-4 stability is reported as half-life at 37 °C (t½, DPP-4, h); plasma pharmacokinetic half-life (t½, pK, h) was determined after a single IV dose (1 mg kg⁻¹) in male ICR mice; GLP-1R potency is EC₅₀ (pM) from a CRE-Luc assay in BHK cells expressing human GLP-1R. Data are mean ± SD (DPP-4 and EC₅₀: three independent experiments; pK: n = 3 mice per group). Statistics are unpaired, two-tailed Welch’s t-tests versus R34-GLP-1: ns, not significant; *P < 0.05; **P < 0.01.

| Peptide | Abbreviation | *In vitro* DPP-4<br>$t_{1/2}$ (h) | *In vitro* receptor activity<br>cAMP assay<br>(EC50, pmol l<sup>-1</sup>) | *In vivo* plasma<br>$t_{1/2}$ (pK) (h) |
| --- | --- | --- | --- | --- |
| [R<sup>34</sup>]-GLP-1(7-37) | R34-GLP-1 | 0.3 ± 0.1 | 11.3 ± 1.2 | 0.08 ± 0.01 |
| [Aib<sup>8</sup>/R<sup>34</sup>]-GLP-1(7-37) | Aib8 (unlipidated<br>semaglutide) | 5.4 ± 0.6** | 8.84 ± 1.2<sup>ns</sup> | 0.32 ± 0.09* |
| [azaAla<sup>8</sup>/R<sup>34</sup>]-GLP-1(7-37) | AzaA8 | >24 | 22.3 ± 2.4** | 0.46 ± 0.12* |
| [azaHis<sup>7</sup>/R<sup>34</sup>]-GLP-1(7-37) | AzaH7 | >24 | 743 ± 164* | 1.67 ± 0.52* |
| [azaHis<sup>7</sup>/Aib<sup>8</sup>/R<sup>34</sup>]-GLP-1(7-37) | AzaH7/Aib8 | >24 | 306 ± 44** | ND |
| [azaHis<sup>7</sup>/azaGlu<sup>9</sup>/R<sup>34</sup>]-GLP-1(7-37) | AzaH7/AzaE9 | >24 | > 10,000 | ND |

Abbreviations/notation: Aib, α-aminoisobutyric acid/ $\alpha$ -methylaniline; aza, α-carbon → nitrogen substitution; R34-GLP-1, Lys34 → Arg variant of GLP-1(7-37); Aib8 (unlipidated semaglutide), Aib<sup>8</sup>/R<sup>34</sup>-GLP-1; AzaA8, azaAla<sup>8</sup>/R<sup>34</sup>-GLP-1; AzaH7, azaHis<sup>7</sup>/R<sup>34</sup>-GLP-1; AzaH7/Aib8, azaHis<sup>7</sup>/Aib<sup>8</sup>/R<sup>34</sup>-GLP-1; AzaH7/AzaE9, azaHis<sup>7</sup>/azaGlu<sup>9</sup>/R<sup>34</sup>-GLP-1. ">24 h" indicates no measurable degradation within the assay window; ND, not detected.

### *In vivo* proteolysis shifts to internal hotspots

To confirm that N-terminal aza-substitution protects the canonical DPP-4 site in a physiological context, we performed tandem mass spectrometry (MS/MS) fragmentation analysis of three analogues (Aib8, AzaA8, and AzaH7) in mice. Peptides were administered intravenously and fragments in whole blood and urine were identified by UPLC–MS/MS over time. All three aza-GLP-1 analogues showed a common pattern of cleavage primarily within the central region of the peptide. A heat map of peptide bond cleavages (**Fig. 2a, b**) revealed recurrent “hotspots” around bonds 12, 14, 17, 19-21, 23-24, and 28-29, corresponding to the sequence FTSDVSSYLEGQAAKEFIAWLVR. In contrast, fragmentation at the extreme N-terminus was greatly reduced. The signature DPP-4 cleavage between positions 7-8 (His-Ala in the native peptide) was notably absent from the dominant fragments. Instead, only low-intensity signals corresponding to the N-terminal dipeptide or its immediate flanking fragments were detected, while high-intensity ions were derived from internal breaks in the peptide (**Fig. 2c–e** and **Supplementary Datasets 1–4**). These data confirm that the aza- (or Aib-) substitution at the His7/Ala8 site effectively blunted cleavage at this locus *in vivo*, causing proteolysis to shift to downstream sites. This site-specific protection suggests that future designs might further improve stability by reinforcing the internal cleavage-prone regions (e.g., around bonds 17–21 and 23–29) identified in the spectra. In summary, MS/MS mapping demonstrates that aza-substitution at the DPP-4 hotspot provides robust protection against N-terminal degradation, with proteolytic breakdown converging on shared internal sites across the different analogues.

**Figure 2.** Cleavage hotspots and dominant whole blood fragments of GLP-1 analogues. **a)** Cleavage hotspots of N-terminus-modified peptides identified by tandem MS experiments. **b)** Peptide bond-specific cleavage intensities are shown for Aib8, AzaA8 and AzaH7. The vertical axis denotes peptide bonds (i|i+1), with circle size and color indicating relative units (RU) of cleavage intensity: red, primary (RU 1.0); orange, secondary (RU 0.5); yellow, low (RU 0.25). All analogues display recurrent hotspots within the central FTSDVSSY…EGQAAKEFIAWLVR region, particularly around bonds 17, 19–21, 23–24, and 28–29. **c-e)** Dominant whole blood fragments for **c)** Aib8, **d)** AzaA8, and **e)** AzaH7, including both the dipeptide and remainder resultant of DPP-4 cleavage, which were consistently detected at very low levels. Collectively, these results show that aza- and Aib-based substitutions reduce N-terminal degradation, but cleavage converges on shared internal hotspots that yield consistent, high-intensity fragments across analogues. Relative units are normalized to the highest value observed for each sample.

### AzaA8 preserves GLP-1 receptor potency

We next assessed whether the aza modifications affected the ability of the peptides to activate the GLP-1 receptor. A cell-based luciferase reporter assay (CRE-Luc) in BHK cells expressing human GLP-1R was used to determine half-maximal effective concentrations (EC_50_) for cAMP signaling (**Table 1**; **Supplementary Table 3**). R34-GLP-1 and Aib8 controls had EC_50_ values of 11.3 ±1.2 pmol l^-1^ and 8.8 ±1.2 pmol l^-1^, respectively, consistent with their high potency. AzaA8 exhibited an EC_50_ of 22.3 ±2.4 pmol l^-1^, indicating that substituting Ala8 with aza-Ala preserved nearly the full agonist potency of the peptide. In contrast, analogues containing an aza-His7 substitution showed somewhat reduced activity: EC_50_ values were in the mid-to-high picomolar or low nanomolar range (e.g., 0.3–0.7 nmol l^-1^ for AzaH7/Aib8 and AzaH7, and >10 nmol l^-1^ for the doubly substituted AzaH7/azaE9). These results identified AzaA8 as the optimal balance of stability and activity, fully resistant to DPP-4 yet retaining low picomolar GLP-1R agonism (**Table 1**).

### Pharmacokinetics reveal extended half-life

We next evaluated how aza-substitutions (AzaA8 and AzaH7) influenced the pharmacokinetic behavior of the peptides *in vivo*. Male CD-1 mice received a single intravenous dose of each peptide (1 mg kg^-1^), and plasma concentrations were monitored over 8 h (**Table 1; Supplementary Table 4**). Unmodified GLP-1(7–37) was rapidly cleared from circulation, with a plasma half-life of approximately 0.08 hours (4.8 minutes), consistent with the peptide’s known instability. Introduction of the Aib substitution at position 8, which is used in semaglutide, extended the plasma half-life to about 0.32 hours (19.2 minutes). Our aza-engineered peptide AzaA8 displayed a further increased half-life to approximately 0.46 hours (27.6 minutes), while AzaH7 persisted for markedly longer, with a half-life of around 1.67 hours (100.2 minutes).

The extended persistence of the aza-analogues in circulation confirms that these substitutions slow peptide clearance *in vivo*. Notably, AzaA8 displayed a longer half-life yet a higher plasma clearance rate (∼43.9 ml min^-1^·kg^-1^) than the Aib8 control (∼22.6 ml min^-1^·kg^-1^). This apparent paradox suggests that although AzaA8 resists enzymatic degradation, it may be removed more rapidly through alternative pathways, such as renal filtration, or may have a reduced volume of distribution. By contrast, AzaH7 showed a markedly lower clearance (∼7.0 ml min^-1^·kg^-1^), consistent with its prolonged circulation time. Overall, incorporation of aza-residues at the peptide N-terminus substantially extended systemic levels relative to native GLP-1, underscoring the stabilizing effect of these modifications.

### Acute glucose lowering in lean mice

To determine whether the enhanced stability of AzaA8 translated into functional efficacy, we tested it in an oral glucose tolerance test (oGTT). Male 21-week-old C57BL/6N mice, maintained on standard chow, were fasted for 6 h with free access to water before the experiment. Groups of weight-matched animals (n = 8 per group) received a single subcutaneous injection at the back of the neck of AzaA8 (0.125 or 0.4 mg kg^-1^), the unlipidated semaglutide analogue Aib8 (0.125 or 0.4 mg kg^-1^), exendin-4 (0.125 mg kg^-1^, positive control), or saline vehicle (**Fig. 3a**). All test articles and controls were blinded to the investigators, and mice were monitored for body weight and clinical condition (**Supplementary Table 5**). Two hours after injection, mice underwent an oral glucose tolerance test: each animal was given a glucose bolus (2 g kg^-1^, delivered as a 100 mg ml^-1^ solution in sterile water) by oral gavage, and blood glucose was measured from tail vein samples at 0, 15, 30, 60, 90, and 120 min.

**Figure 3.** Effect of acute administration of AzaA8 versus controls on oral glucose tolerance test (oGTT) in lean (chow-fed) C57BL/6N mice. **b)** Blood glucose levels over time (min) during oGTT, and **c)** total average area under curve for oGTT blood glucose levels – in male 21-week-old, chow-fed lean mice (C57Bl/6N) after single subcutaneous injection (treatment -2 h) of GLP-1 analogues or vehicle (saline) prior to glucose bolus (t = 0), n = 8 animals per group. Data are presented as mean ± SEM. Statistical analysis in **b** was performed using two-way ANOVA followed by Dunnett’s multiple comparisons test. P-values are indicated in the graphs and ****p < 0.0001, comparing each test article group to vehicle. When p-value is not indicated the comparison was not significant. Statistical significance in **c** was performed by one-way ANOVA followed by Tukey’s multiple comparisons test (all pairwise). P-values are indicated in the graphs and ****p < 0.0001. Panel **a** created with BioRender.com.

AzaA8 significantly reduced glycemic excursion in a dose-dependent manner (**Fig. 3b, c**, **Supplementary Fig. 1, and Supplementary Table 6**). At the higher dose of 0.4 mg kg^-1^, AzaA8 nearly matched the glucose-lowering efficacy of Aib8 and approached that of exendin-4. Even at 0.125 mg kg^-1^, AzaA8 blunted the glucose rise relative to saline-treated controls. These findings demonstrate that the stability conferred by aza-substitution translates into preserved biological efficacy *in vivo*, enabling AzaA8 to provide acute glycemic control comparable to clinically validated GLP-1 receptor agonists like Exendin-4.

### Sustained metabolic benefits in obese mice

Encouraged by the acute results, we evaluated whether AzaA8 could provide longer-term metabolic benefits in animals with high-fat diet-induced obesity (DIO). Male C57BL/6N mice were fed a 60% fat diet from 9 weeks of age until they became obese (body weight ≥40 g at ∼15-21-week old; see **Fig. 4a**), then allocated to treatment groups (n = 7-8 per group) for a 4-week intervention. AzaA8 was administered at 0.4 mg kg^-1^ day^-1^ (split into twice-daily intraperitoneal doses), and its effects were compared to daily liraglutide (0.4 mg kg^-1^, i.p., twice daily as described above), intermittent semaglutide (0.12 mg kg^-1^, i.p., every 3 days), or saline vehicle.

**Figure 4.** Effect of chronic intraperitoneal (i.p.) administration of GLP-1 receptor agonists on body weight and glucose levels in high-fat diet-induced obesity (HF-DIO) model. **a)** The model consists in high-fat diet (60%)-fed C57BL/6N male mice (at least 15-weeks-old, or ≥40 g) treated for 29 days, **b)** bi-weekly body weights and, **c)** total body weight loss for AzaA8 (0.4 mg kg^-1^ day^-1^, BID, every 12 h, liraglutide (0.4 mg kg^-1^ day^-1^, BID, every 12 h), semaglutide (0.12 mg kg^-1^ 3-day^-1^, QD) or vehicle (saline BID, every 12 h), **d)** blood glucose levels after oral glucose tolerance test (treatment day 29) in HF-DIO mice treated daily as indicated for n = 7-8 animals per group. Effect of chronic (i.p.) administration of lead GLP-1R agonist-AzaA8 *vs.* controls on blood levels of selected diabetes- and metabolism-related proteins in HF-DIO model. Fasting **e)** leptin, **f)** insulin, **g)** glucagon, **h)** total PYY, and **i)** HbA1c levels in in C57BL/6N DIOmice after 29 days of treatment with vehicle (daily), 0.12 mg kg^-1^ 3-day^-1^ semaglutide, 0.4mg kg^-1^ day^-1^ liraglutide or AzaA8 (n = 7-8 mice per group). Effect of chronic (i.p.) administration of GLP-1 receptor agonists on isolated tissue weights in HF-DIO model. **j)** iWAT, **k)** iBAT, **l)** quadriceps skeletal muscle, and **m)** liver wet masses in C57BL/6N DIO mice after 29 days of daily treatment with vehicle (daily), 0.12 mg kg^-1^ 3-day^-1^ semaglutide, 0.4 mg kg^-1^ day^-1^ liraglutide or AzaA8 (n = 7-8 mice per group). Data are presented as mean ± SEM unless otherwise indicated. For panels **b** and **d**, group differences were analyzed using two-way ANOVA followed by Dunnett’s multiple comparisons test (vehicle vs. test article groups). For panel **c**, one-way ANOVA followed by Tukey’s multiple comparisons test (all pairwise) was used. For panels **e**-**i**, data are shown as mean ± SD and analyzed by one-way ANOVA with Dunnett’s multiple comparisons test (vehicle vs. test article groups). For panels **j**-**m**, data are shown as mean ± SEM and analyzed by one-way ANOVA with Dunnett’s multiple comparisons test (vehicle vs. test article groups). P-values are indicated in the graphs and ****p < 0.0001, when p-value is not indicated the comparison was not significant. Panel **a** created with BioRender.com.

AzaA8-treated mice exhibited a significant reduction in body weight over the 29-day treatment, losing on average ∼16% of their starting weight (**Fig. 4b, c**). This degree of weight loss was comparable to that achieved by liraglutide (∼21.5%) at the same daily dose (∼0.4 mg kg^-1^) and was slightly greater than the weight reduction observed with the unlipidated semaglutide peptide (∼9.5%, Aib8, 0.4 mg kg^-1^ day^-1^) (**Supplementary Fig. 2**). Semaglutide (administered at 0.12 mg kg^-1^ every 3 days) also induced substantial weight loss (∼19.8% in this model), as expected^14^. Thus, despite lacking the fatty-acid prolongation motif of semaglutide, AzaA8 delivered robust metabolic efficacy when given daily.

Glucose control was likewise improved by AzaA8. At the end of the 4-week treatment, mice were subjected to an oGTT. AzaA8 significantly blunted the glucose rise compared to vehicle-treated DIO mice (**Fig. 4d**). The glucose excursion in the AzaA8 group was similar to that in the semaglutide-treated group and markedly lower than in the vehicle group. Interestingly, liraglutide-treated mice did not show a significant improvement in oGTT glucose levels in this experiment, despite their weight loss. This contrasts with the typical profile of liraglutide and may relate to the specific dosing schedule or model variability^15,16^.

To further characterize metabolic changes, we measured key hormones and biomarkers at the end of the study. Fasting serum leptin (**Fig. 4e**) and insulin (**Fig. 4f**) levels were both significantly reduced in the AzaA8 group relative to vehicle, consistent with a loss of adipose mass and improved insulin sensitivity. Meanwhile, fasting glucagon (**Fig. 4g**) and peptide tyrosine tyrosine (PYY) levels (**Fig. 4h**) were unchanged, and glycated hemoglobin (HbA1c) remained low (∼4.5–4.9%) and similar across all groups (**Fig. 4i** and **Supplementary Table 7**). The reduction in leptin in AzaA8-treated mice likely reflects decreased fat mass, as suggested by direct measurements of adipose tissue. AzaA8 significantly reduced the weight of subcutaneous inguinal white adipose tissue (iWAT) compared to vehicle (1220.8 ± 232.9 mg vs 1993.8 ± 328.8 mg, **Fig. 4j**). This reduction was comparable to that seen with semaglutide (1206.1 ± 340.9 mg) and liraglutide (907.9 ± 279.4 mg). Importantly, AzaA8 had minimal effect on the mass of interscapular brown adipose tissue (iBAT) (**Fig. 4k**) and did not cause loss of lean tissue such as skeletal muscle (quadriceps) (**Fig. 4l**) or significant liver weight change (**Fig. 4m**). Preservation of iBAT and muscle, alongside preferential loss of white fat, is a favorable outcome, suggesting that weight loss occurred mainly through depletion of excess white adipose tissue (**Supplementary Fig. 3)**. Finally, to assess safety, we profiled inflammatory cytokines in the serum. We observed no significant changes in IL-6, IL-1β, TNF-α, or other cytokines in AzaA8-treated mice versus controls (**Supplementary Fig. 4**), indicating that the 4-week treatment did not provoke an adverse inflammatory response.

To test the clinically relevant subcutaneous route of administration (**Fig. 5a**), we performed a parallel four-week study in diet-induced obese mice. Daily injections of AzaA8 (0.4 mg kg^-1^, s.c., daily), stabilized body weight relative to vehicle, preventing further gain, while semaglutide (0.04 mg kg^-1^, s.c., daily) produced modest weight loss (**Fig. 5b, c**). Food intake was slightly reduced in the AzaA8 group during the latter half of the treatment period (**Fig. 5d** and **Supplementary Table 8**).

**Figure 5.** Effect of chronic subcutaneous (s.c.) administration of GLP-1 receptor agonists on body weight and glucose levels in HF-DIO model. **a)** High-fat diet (60%)-fed C57BL/6N male mice treated for 29 days with s.c. administration, **b)** bi-weekly body weights, **c)** total body weight loss, **d)** food consumption for AzaA8 (0.4 mg kg^-1^ day^-1^), semaglutide (0.04 mg kg^-1^ day^-1^) or vehicle (saline), **e)** blood glucose levels after oral glucose tolerance test (treatment day 29), and **f)** total glucose AUC in HF-DIO mice treated daily as indicated for n = 8 animals per group; Data are presented as mean ± SEM unless otherwise indicated. In panels **b**, **d** and **e**, group differences were analyzed using two-way ANOVA followed by Dunnett’s multiple comparisons test (vehicle vs. test article groups). In panels **c** and **f**, group differences were analyzed by one-way ANOVA followed by Tukey’s multiple comparisons test. P-values are indicated in the graphs and ****p < 0.0001, when p-value is not indicated the comparison was not significant. Panel **a** created with BioRender.com.

Glycemic control outcomes mirrored those of the i.p. study. At the end of 4 weeks, AzaA8-treated mice exhibited significantly improved glucose tolerance in the oGTT, with glucose levels during the test being the lowest among the groups (**Fig. 5e**). The total glucose AUC was significantly reduced by AzaA8 (764.7±81.5 mmol l^-1^ min^-1^) compared to vehicle (1742.5±104.9 mmol l^-1^ min^-1^) and even somewhat lower than in the semaglutide group (1195.4±89.2 mmol l^-1^ min^-1^) (**Fig. 5f**). Daily monitoring of non-fasting blood glucose provided additional insight: when measured 2 h after dosing, AzaA8-treated mice showed acutely lower glucose levels relative to vehicle (**Supplementary Fig. 5a**), whereas measurements taken just before each dose (pre-dose baseline glucose levels) showed no significant differences between groups (**Supplementary Fig. 5b**). This indicates that AzaA8’s glucose-lowering effect was closely tied to its dosing/exposure and that the HFD-fed mice in this timeframe did not develop overt fasting hyperglycemia. Consistent with that the above, HbA1c remained in the normal range (∼3.6–3.8%) for all groups and did not differ significantly (**Supplementary Table 9**), reflecting that this DIO model, while obese and insulin-resistant, had not progressed to frank diabetes within the study duration. Overall, these chronic studies demonstrate that AzaA8 delivers sustained metabolic benefits, including weight reduction, preferential fat loss, improved glucose tolerance, and favorable hormonal changes, without triggering inflammation. The efficacy of AzaA8 across both intraperitoneal and subcutaneous routes of adminstration highlights the robustness of its therapeutic potential.

## Discussion

Peptidomimetics with minimal backbone modifications can retain efficacy of native function while gaining important pharmacological advantages. In this study, we demonstrate that a single α-aza-amino acid substitution at a critical proteolytic site endows a GLP-1 analogue with markedly improved stability and *in vivo* efficacy. Replacing Ala8 with aza-alanine “locks” the N-terminus against DPP-4 attack, extending the peptide’s half-life and enabling sustained receptor agonism *in vivo*. This design leaves the peptide’s sequence and structure virtually unchanged aside from one atom, thus minimizing disruption of GLP-1R binding and likely reducing immunogenic risk^17,18^.

Our results show that AzaA8 performs on par with established and clinically approved GLP-1RAs in animal models: it improved glucose tolerance acutely and produced significant weight loss or weight stabilization with chronic dosing, comparable to the effects of liraglutide and semaglutide. Notably, AzaA8 achieved these benefits without the need for fatty acid derivatization or other extensive modifications beyond the aza-substitution. The differences observed between our intraperitoneal and subcutaneous dosing studies (e.g. a greater magnitude of weight loss in the i.p. regimen) likely reflect differences in pharmacokinetics (exposure levels) and dosing frequency, as well as the comparator doses used. Nonetheless, both studies consistently highlight AzaA8’s potent metabolic effects and support its efficacy across different administration routes.

Our findings align with a recent report by Dinsmore *et al*.^19^, who independently identified that an aza-Ala8 substitution in GLP-1 analogues confers resistance to proteolysis and prolongs peptide half-life *in vitro* while retaining agonist activity. While Dinsmore *et al.* focused on *in vitro* stability and receptor assays, our work extends the evidence to *in vivo* efficacy, showing that an aza-GLP-1 analogue can deliver meaningful therapeutic benefits in animal models of diabetes and obesity. The convergence of these results from separate groups strengthens the case for aza-amino acids as valuable tools in peptide drug design. Collectively, these findings position aza-substitution as a rational and intrinsically efficient strategy to stabilize therapeutic peptides while preserving native function—marking a shift from empirical modification toward data-driven, atom-level peptide design that could simplify manufacturing and reduce development costs for durable peptide therapeutics.In summary, targeted aza modifications represent a promising approach to bolster peptide therapeutics. By blocking a primary proteolytic hotspot, we created a GLP-1 analogue that is both long-lasting and highly active, achieving significant anti-diabetic and anti-obesity effects in mice. This proteolytic-hotspot aza-subsitution strategy should be generalizable to other peptide hormones or therapeutics that suffer from rapid degradation, opening the door to more durable and effective peptide-based treatments for metabolic disease and beyond.

## Methods

### Reagents and chemicals

All chemicals and reagents were purchased from Fisher Scientific (Hampton, NH) or from Sigma-Aldrich (St. Louis, MO) unless otherwise noted. GLP-1 (7-37) was obtained from Tocris Bioscience™ (Minneapolis, MN), while semaglutide and exendin-4 were obtained from MedChemExpress LLC (Monmouth Junction, NJ).

### Azapeptide analogues and peptide synthesis

Peptide controls and azapeptides were synthesized using standard reagents and methods on a Liberty Blue™ 2.0 Microwave Peptide Synthesizer (CEM Corporation, Matthews, NC). Aza-amino acids and azaGLP-1 analogues were synthesized using our previously established azapeptide synthesis platform, employing benzotriazole-based building blocks and microwave-assisted SPPS^13^.

### Structure verification and purity methods

Flash Chromatography and HPLC were used to purify all synthesized peptides and aza-analogues (purity level minimum >95%), and high-resolution mass spectrometry instrument (HRMS) was used to confirm sequence/structures (Agilent 6550 iFunnel QTOF LC/MS). Detailed support for characterization, structural confirmation and purities of the GLP-1 azapeptide analogues are provided in^13^.

### *In vitro* protease stability: DPP-4 half-life assay

Stock solutions (5 mg ml^-1^) of test articles and positive controls were made in sterile ddH_2_O. Reactions were prepared by incubating 16 nmol l^-1^ DPP-4 (D3446, Aldrich) at 37 °C, with a final test article concentration of 0.1 mmol l^-1^ in sterile Tris HCl buffer (50 mmol l^-1^, pH 7.4). The vials were kept in a 37 °C incubator for the duration of the experiment. Aliquots (30 μl) were taken at various timepoints (0, 1, 4, 8, 24 h) and added to vials which had been pre-filled with 20 μl of 0.05% TFA in water, and gently mixed. The samples were injected directly into the HPLC to monitor the recovery of intact peptide. The HPLC (Waters (Breeze QS System) was equipped with a 1525 binary pump, and a Phenomenex kinetex 2.6 mm EVO C18 analytical column (100 Å 150 × 4.6mm). Chromatography was performed at ambient temperature with flow rate of 0.7 ml min^-1^ with linear gradient from water (0.05% TFA): MeOH (0.05% TFA) [95:5] to water (0.05% TFA): MeOH (0.05% TFA) [5:95] over 15 min and resolved peaks were detected by a 2998 photodiode array (PDA) detector at 215 nm. The peak area response ratio (PARR) was compared to the PARR at time 0 to determine the percentage of test article remaining at each time point. Half-lives of samples in DPP-4 were calculated using Microsoft Excel. The general methods were adapted from^20^.

### UPLC–MS/MS profiling of N-terminally modified GLP-1 analogues

Saline solutions (0.2 mg ml^-1^) of each test compound were administered to male ICR mice (n = 2) via a single intravenous injection (1 mg kg^-1^). Blood and urine samples (100–200 µL) were collected at 0, 15, and 60 min post-dose and immediately quenched in pre-chilled tubes containing 200–400 µL of acetonitrile:methanol (1:1, v/v) to precipitate proteins and halt proteolysis. Samples were vortexed and held on ice (∼4 °C) until the final time point, then vortexed again and centrifuged at 18,200 ×g for 10 min. Supernatants were removed, diluted 1:1 with 2% acetonitrile in 0.1% formic acid (water), filtered, and analyzed by HPLC-MS/MS; when not injected immediately, extracts were maintained on dry ice (−78 °C) prior to analysis. Cleavage site mapping was carried out on a Waters ACQUITY UPLC–TQD with PDA detection (190-400 nm). Peptides were resolved on an ACQUITY UPLC HSS C_18_ column (1.8 µm, 2.1 × 50 mm) with mobile phase A (water + 0.1% formic acid) and B (acetonitrile + 0.1% formic acid), 2 µl injections, 0.3 ml min^-1^. MS was acquired by ESI^+^ with full-scan MS^1^ (m/z 200-2,000) and product-ion MS/MS (m/z 100-2,000) using collision-energy ramps (e.g., 15-35 eV). Targeted transitions/precursors for each peptide (e.g., [M+3H]^3+^, [M+4H]^4+^) were subjected to product-ion scans, and b/y ion series were assigned with a 10 ppm tolerance.

### *In vivo* stability-murine pharmacokinetic (pK) studies

A plasma pK study (Pharmacology Discovery Services Taiwan, Ltd.) was performed in male ICR mice (n = 6) following single intravenous (IV) administration of test articles at 1 mg kg^-1^. The semi-serial plasma samples were collected from three alternative animals at each time point at 0.05, 0.167, 0.5, 1, 2, 4, 6 and 8 h after IV administration. The body weight of each animal was recorded and exposure levels (ng ml^-1^) of test articles in plasma samples determined by liquid chromatography–tandem mass spectrometry (LC-MS/MS^21^). The exposure levels (ng ml^-1^) of test articles in plasma samples were determined by LC-MS/MS. Results were plotted for plasma concentrations versus time (mean ± SD). The fundamental pK parameters after IV administration were obtained from the NCA of the plasma data using WinNonlin (manual mode). The lower limit values of quantification (LLOQ) in plasma samples was 2 ng ml^-1^, and the exposure levels below the 75% of LLOQ (1.5 ng ml^-1^) were determined as below the limit of quantification (BLOQ). The fundamental pK parameters of test articles after IV (t_1/2_, C_0_, AUC_last_, AUC_inf_, AUC/D, AUC_extr_, MRT, Vss and CL) administration were obtained from the non-compartmental analysis (NCA) of the plasma data using WinNonlin.

### *In vitro* GLP-1 receptor potency measured by cAMP signaling

The *in vitro* potency assay (Novo Nordisk) relied on a reporter gene assay (CRE-luc) and was carried out in baby hamster kidney (BHK-21) cells overexpressing the hGLP-1 receptor. For these experiments, the compounds were tested in four technical pseudo-replicates. The combined results from three independent experiments were used to determine EC_50_ values and data analyzed for significance at a 95% confidence interval.

### Acute efficacy of AzaA8 in oral glucose tolerance test (oGTT) in lean mice

*In vivo* studies were conducted at Pharmaron (Beijing, China) in compliance with Institutional Animal Care and Use Committee (IACUC) protocols and Association for Assessment and Accreditation of Laboratory Animal Care (AAALAC) guidelines. Based on the statistical power calculations, eight animals per group were used. Male C57BL/6N mice (21-week old, chow fed; SPF Beijing Laboratory Animal Technology Co., Ltd) were fasted for 6 h with free access to water before testing. At 2 h prior to oGTT, baseline blood glucose was measured, followed by a single subcutaneous injection of the test articles, AzaA8 (0.125 or 0.4 mg kg^-1^), Aib8 (0.125 or 0.4 mg kg^-1^), exendin-4 (0.125 mg kg^-1^), or saline. Treatments were blinded to the investigators, and mice were monitored for clinical abnormalities (**Supplemental Table 5**). For the oGTT, body weights were recorded and at the start of the experiment mice were given 2 g kg^-1^ of glucose (100 mg ml^-1^) by oral gavage, and blood glucose was monitored at 0, 15, 30, 60, 90, and 120 min using an Accu-Chek Guide glucometer from tail blood.

### Chronic *in vivo* efficacy in obese mice

Chronic efficacy studies were conducted at Pharmaron (Beijing, China) under IACUC- and AAALAC-approved protocols, designed to parallel the acute study. Male C57BL/6N mice (SPF Beijing Laboratory Animal Technology Co., Ltd) were placed on a high-fat diet (HFD; D12492, Research Diets, Inc.) at 7 weeks of age and maintained for 15 weeks before allocation into weight-matched groups (n = 8). Mice were monitored for clinical abnormalities (**Supplemental Table 5**).

For subcutaneous dosing, animals (18–21 weeks old) received daily treatment for 4 weeks: AzaA8 (125 µg kg^-1^ during week 1, then 0.4 mg kg^-1^ in weeks 2-4; once daily in week 2, twice daily from week 3 onward), semaglutide (0.04 mg kg^-1^ daily), or saline vehicle. Treatments were blinded to investigators, and mice were monitored daily for health. Body weight, food intake, and non-fasting blood glucose were measured twice weekly (2 h post-dose and immediately before dosing; **Supplementary Fig. 5**). At study end, mice underwent an oral glucose tolerance test (oGTT) following a 6 h fast. Blood glucose was recorded before dosing (-2 h), test compounds administered, and a 2 g kg^-1^ glucose challenge delivered by oral gavage at time zero. Glucose levels were monitored at 0, 15, 30, 60, 90, and 120 min using a handheld glucometer. Mice were euthanized after 120 min, and blood was collected for HbA1c analysis.

For intraperitoneal dosing, studies were performed at the Feinstein Institutes (protocol #24-0467). Male C57BL/6N mice (Taconic Biosciences, USA) were fed HFD (D12492) from 9 weeks of age for ≥10 weeks before treatment initiation (≥15-week old, ≥40 g). Animals (n = 8 per group) received AzaA8 or liraglutide (0.4 mg kg^-1^ twice daily, i.p.), semaglutide (0.12 mg kg^-1^ every 3 days), or saline vehicle for 4 weeks. Body weight was recorded twice weekly. At study completion, mice underwent oGTT as above, with glucose monitored up to 120 min using a Vet GlucoGauge (Covetrus). Blood and serum were collected for HbA1c (Mouse Hemoglobin A1c (HbA1c) Kits (Catalog# 80310, Crystal Chemistry, Elk Grove Village, IL)), cytokine profiling (IFN-γ, IL-1β, IL-6, IL-10, IL-12p70, KC/GRO, TNF-α using an MSD kit (Proinflammatory Panel 1 Mouse Kit, 7 plex, K15048G-1, Meso Scale Diagnostics, LLC (MSD), Rockville, MD), and metabolic hormone measurements (insulin, leptin, glucagon, PYY; using an MSD kit (U-PLEX Custom Metabolic Group 1 Assays, K152ACM-1, MSD, Rockville, MD)). Tissue depots (iWAT, iBAT, quadriceps muscle, liver) were excised, weighed, and frozen for further analysis.

## Reporting summary

Further information on research design is available in the Nature Portfolio Reporting Summary linked to this article.

## Data availability

Source data are provided with this paper.

## Code availability

This work did not generate code.

## Acknowledgments

We would like to thank Dr. Michael Brines for critical reading of the manuscript. Molecules were rendered using the PyMOL Molecular Graphics System, Version 3.1.1 Schrödinger, LLC. This work was supported by funding from the Feinstein Institutes for Medical Research. Some of this work was supported by Northwell Health’s 2019 Innovation Challenge prize. Research reported in this preprint was supported by the National Institute of General Medical Sciences of the National Institutes of Health under award number R35GM138201 and by the Defense Threat Reduction Agency under award number HDTRA1-21-1-0014. The content is solely the responsibility of the authors and does not necessarily represent the official views of the National Institutes of Health or the Defense Threat Reduction Agency.

This preprint has not been peer-reviewed and should not be used to guide clinical practice.

## Authors contributions

M.H., S.V.P, and Y.A. designed the study; M.H., K.F.C. synthesized and characterized the peptides. M.H., B.A., and I.T.M., ran animal experiments and analyzed data. B.B. performed and analyzed cAMP signaling and GLP-1 receptor binding assay. M.S. performed organ and tissue collection, serum cytokine and metabolic analyses, and interpreted the resulting data. M.D.T.T. and C.d.l.F.-N. designed, performed, analyzed the *in vivo* proteolysis experiments. M.H., S.V.P., M.D.T.T., C.d.l.F.-N., and Y.A., wrote the manuscript. All authors contributed to manuscript revision and editing.

## Supplemental Data and Figures

**Supplementary Table 1.**
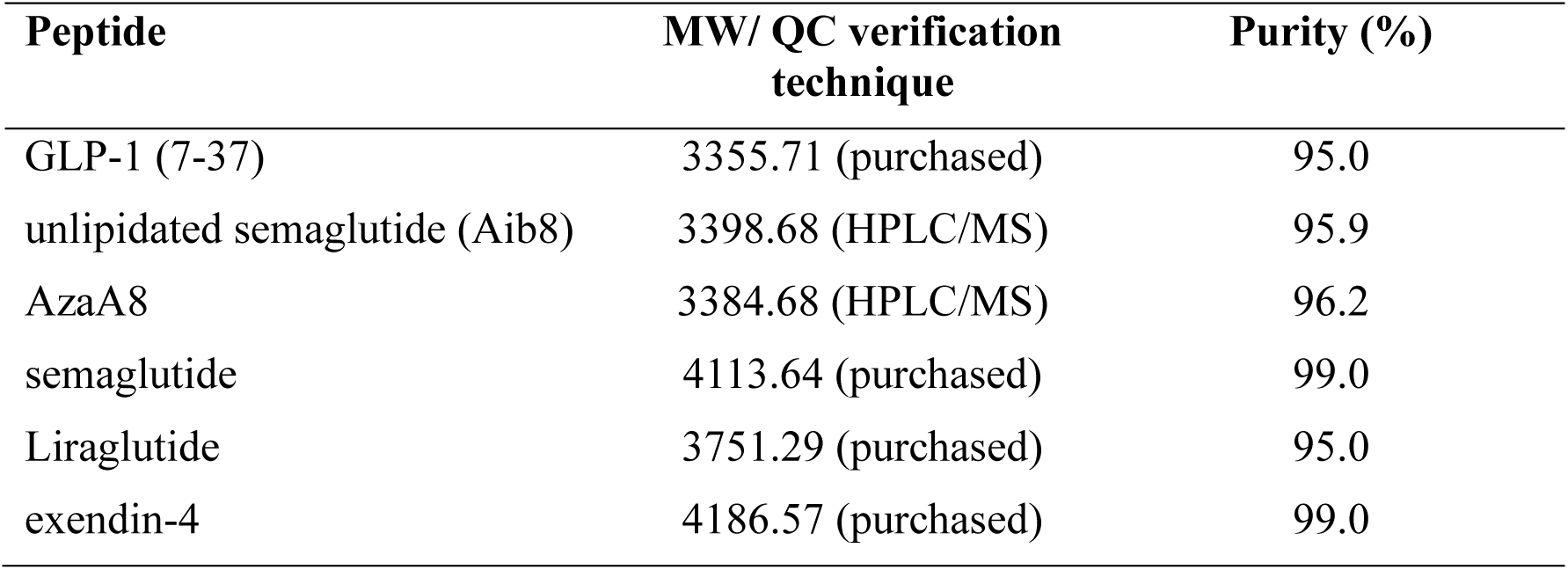
Summary of peptide identities and purity (from Reference 13). Each peptide’s molecular weight was confirmed by MS, and purity was determined by analytical HPLC.

**Supplementary Table 2.** *In vitro* DPP-4 stability of GLP-1 analogues. Values indicate the percentage of intact peptide remaining at each time point during incubation with DPP-4, and the calculated half-life (t½). “>24 h” indicates no measurable degradation within 24 h.

| Aza-GLP-1 analogues | Remaining peptide (%) | | | | | *In vitro* DPP-4 stability ($t_{1/2}$; h) |
| --- | --- | --- | --- | --- | --- | --- |
|  | t = 0 | t = 1 h | t = 4 h | t = 8 h | t = 24 h |  |
| [R<sup>34</sup>]-GLP-1(7-37) | 100 | 9.2 | <5 | - | - | 0.3 ± 0.1 |
| [Aib<sup>8</sup>/R<sup>34</sup>]-GLP-1(7-37) | 100 | 76.2 | 72.3 | 40.3 | 20.6 | 5.4 ± 0.6 |
| [azaAla<sup>8</sup>/R<sup>34</sup>]-GLP-1(7-37) | 100 | 100 | 100 | 100 | >95 | >24 |
| [azaHis<sup>7</sup>/R<sup>34</sup>]-GLP-1(7-37) | 100 | 100 | 100 | 100 | >96 | >24 |
| [azaHis<sup>7</sup>/Aib<sup>8</sup>/R<sup>34</sup>]-GLP-1(7-37) | 100 | 100 | 100 | 100 | >96 | >24 |
| [azaHis<sup>7</sup>/azaGlu<sup>9</sup>/R<sup>34</sup>]-GLP-1(7-37) | 100 | 100 | 100 | 100 | >95 | >24 |

**Supplementary Table 3.** Human GLP-1 receptor cAMP (Cre-Luc) activation assay. Representative dose-response curves for various GLP-1(7–37) analogues were generated using a cAMP-response element luciferase (Cre-Luc) reporter system to assess activation of the human GLP-1 receptor. EC₅₀ values were calculated from these concentration–response curves and are summarized in the table below. Data were provided by Novo Nordisk (Måløv, Denmark). Among the tested peptides, [Aib⁸/R³⁴]-GLP-1(7–37) showed the highest potency, followed by [R³⁴]-GLP-1(7–37) and [azaAla⁸/R³⁴]-GLP-1(7–37), while modifications at position 7 and 9 (azaHis⁷ and azaGlu⁹) significantly reduced receptor activation potency.

| Compound | Mean EC<sub>50</sub> (95% CI), [M] |
| --- | --- |
| [R<sup>34</sup>]-GLP-1(7-37) | 1.13E-11 (8.58E-12 to 1.48E-11) |
| [Aib<sup>8</sup>/R<sup>34</sup>]-GLP-1(7-37) | 8.84E-12 (6.43E-12 to 1.20E-11) |
| [azaAla<sup>8</sup>/R<sup>34</sup>]-GLP-1(7-37) | 2.23E-11 (1.73E-11 to 2.85E-11) |
| [azaHis<sup>7</sup>/R<sup>34</sup>]-GLP-1(7-37) | 7.43E-10 (4.01E-10 to 1.32E-09) |
| [azaHis<sup>7</sup>/Aib<sup>8</sup>/R<sup>34</sup>]-GLP-1(7-37) | 3.06E-10 (2.11E-10 to 4.35E-10) |
| [azaHis<sup>7</sup>/azaGlu<sup>9</sup>/R<sup>34</sup>]-GLP-1(7-37) | > 1E-8 |

**Supplementary Table 4.**
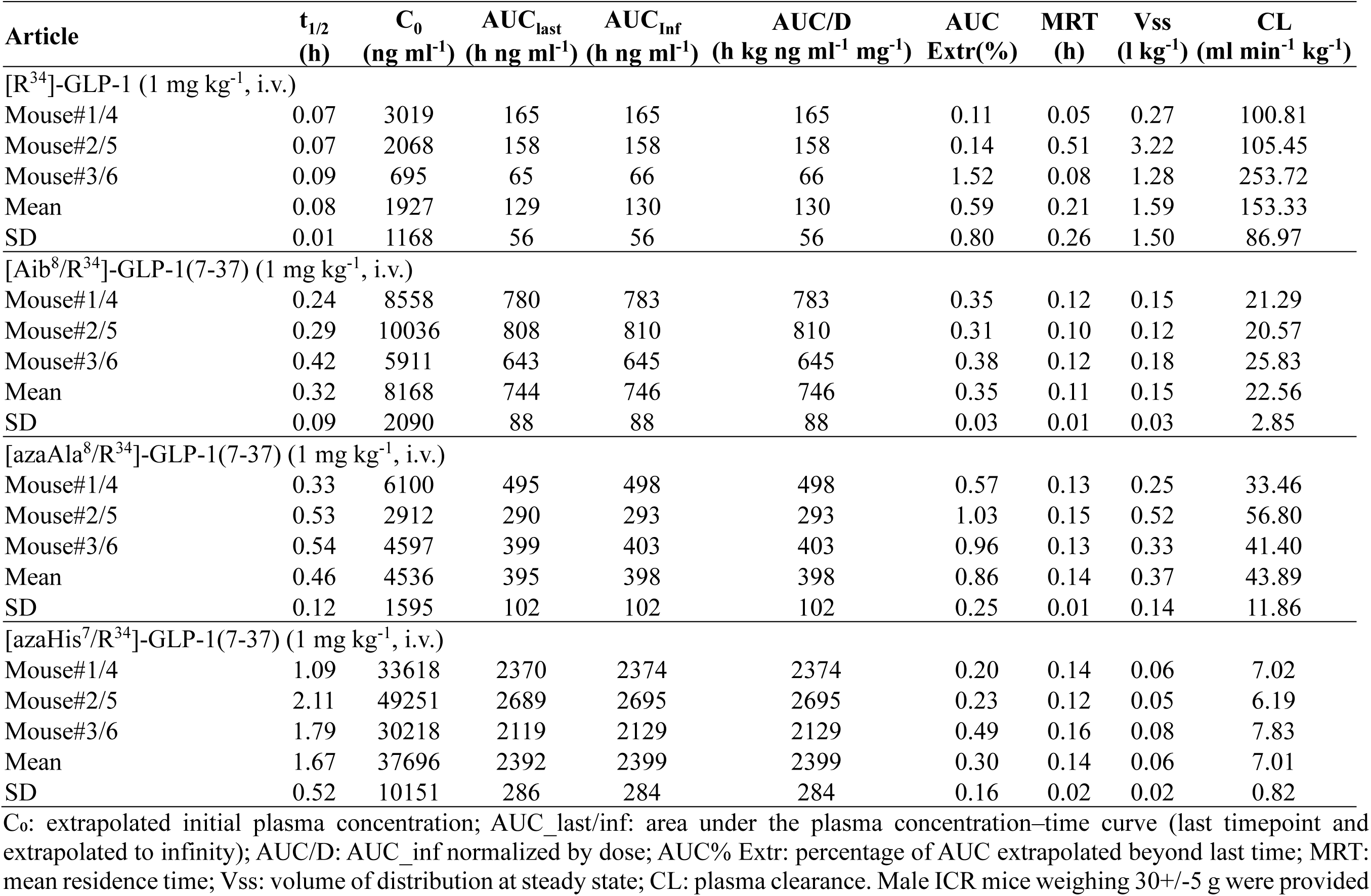
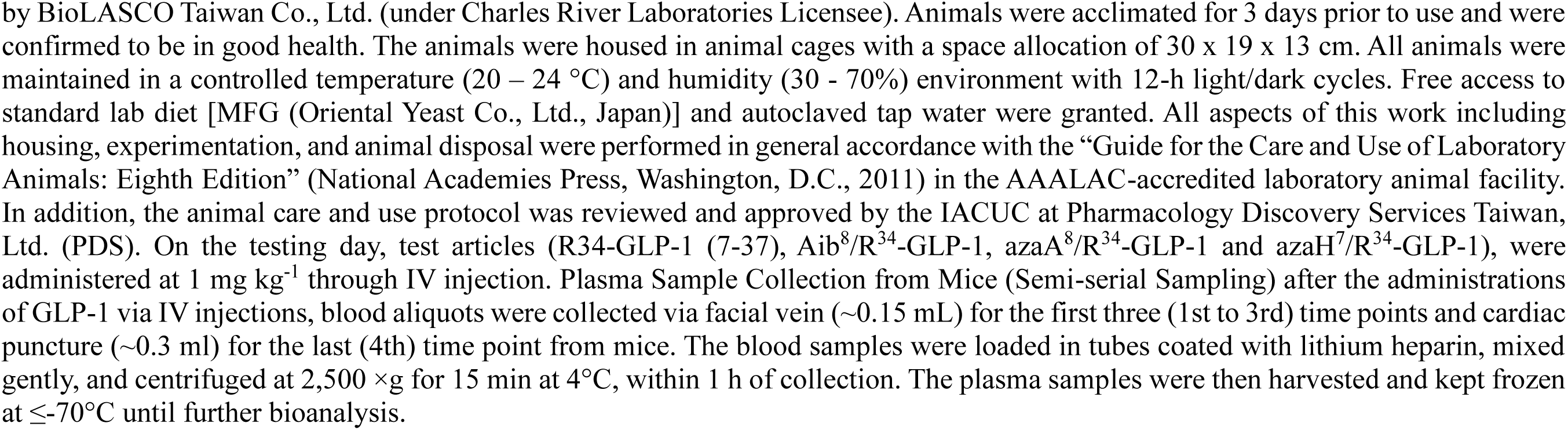
Pharmacokinetic parameters of GLP-1 analogues in male ICR mice (1 mg/kg IV bolus). Data are individual values for three mice per group (semi-serial sampling), with the mean ±S.D. shown below each group. C₀: extrapolated initial plasma concentration; AUC_last/inf: area under the plasma concentration–time curve (last timepoint and extrapolated to infinity); AUC/D: AUC_inf normalized by dose; AUC% Extr: percentage of AUC extrapolated beyond last time; MRT: mean residence time; Vss: volume of distribution at steady state; CL: plasma clearance. Male ICR mice weighing 30+/-5 g were provided by BioLASCO Taiwan Co., Ltd. (under Charles River Laboratories Licensee). Animals were acclimated for 3 days prior to use and were confirmed to be in good health. The animals were housed in animal cages with a space allocation of 30 x 19 x 13 cm. All animals were maintained in a controlled temperature (20 – 24 °C) and humidity (30 - 70%) environment with 12-h light/dark cycles. Free access to standard lab diet [MFG (Oriental Yeast Co., Ltd., Japan)] and autoclaved tap water were granted. All aspects of this work including housing, experimentation, and animal disposal were performed in general accordance with the “Guide for the Care and Use of Laboratory Animals: Eighth Edition” (National Academies Press, Washington, D.C., 2011) in the AAALAC-accredited laboratory animal facility. In addition, the animal care and use protocol was reviewed and approved by the IACUC at Pharmacology Discovery Services Taiwan, Ltd. (PDS). On the testing day, test articles (R34-GLP-1 (7-37), Aib^8^/R^34^-GLP-1, azaA^8^/R^34^-GLP-1 and azaH^7^/R^34^-GLP-1), were administered at 1 mg kg^-1^ through IV injection. Plasma Sample Collection from Mice (Semi-serial Sampling) after the administrations of GLP-1 via IV injections, blood aliquots were collected via facial vein (∼0.15 mL) for the first three (1st to 3rd) time points and cardiac puncture (∼0.3 ml) for the last (4th) time point from mice. The blood samples were loaded in tubes coated with lithium heparin, mixed gently, and centrifuged at 2,500 ×g for 15 min at 4°C, within 1 h of collection. The plasma samples were then harvested and kept frozen at ≤-70°C until further bioanalysis.

## Supplementary Methods. Pharmaron Animal Maintenance protocols

· 1.2.1 Quarantine: Animals were quarantined for at least 14 days before the study. The general health of the animals was evaluated by a veterinarian, and complete health checks were performed. Animals with abnormalities were excluded prior the study.

· 1.2.2 Housing: General procedures for animal care and housing were in accordance with the standard, Commission on Life Sciences, National Research Council, Standard operating procedures (SOPs) of Pharmaron, Inc. The mice were kept in laminar flow rooms at constant temperature and humidity with three animals in each cage. Animals were housed in polycarbonate cages and in an environmentally monitored, well-ventilated room maintained at a temperature of (22 ± 3℃) and a relative humidity of 40 %-80 %. Fluorescent lighting provided illumination approximately 12 hours per day. The bedding material were soft wood, which were changed once per week. Analysis of bedding contamination were conducted annually per facility SOPs and the results of the analyses were retained in the facility records.

· 1.2.3 Animal ID: Each animal was assigned an identification number and marked in the tail; the following identification method was applied. Each cage card was labeled with such information as study number, group, sex, dose, animal number, initiation date, study director.

· 1.2.4 Diet: All animals have free access to standard chow (single dose, lean animals study) or HFD diet (D12492) (chronic doses, HF-DIO study) during the entire study period except for time periods specified by the oGTT protocol (pre-test 6hr fast).

· 1.2.5 Water: Sterile drinking water in a bottle was available to all animals’ ad libitum during the quarantine and study periods. The bottle and the stopper with attached sipper tube were autoclaved prior to use. Samples of water from the animal facility were analyzed and results of water analysis were retained in the facility records and were reviewed by the veterinarian, or designee, to assure that no known contaminants were present that could interfere with or affect the outcome of studies.

**Supplementary Table 5.**
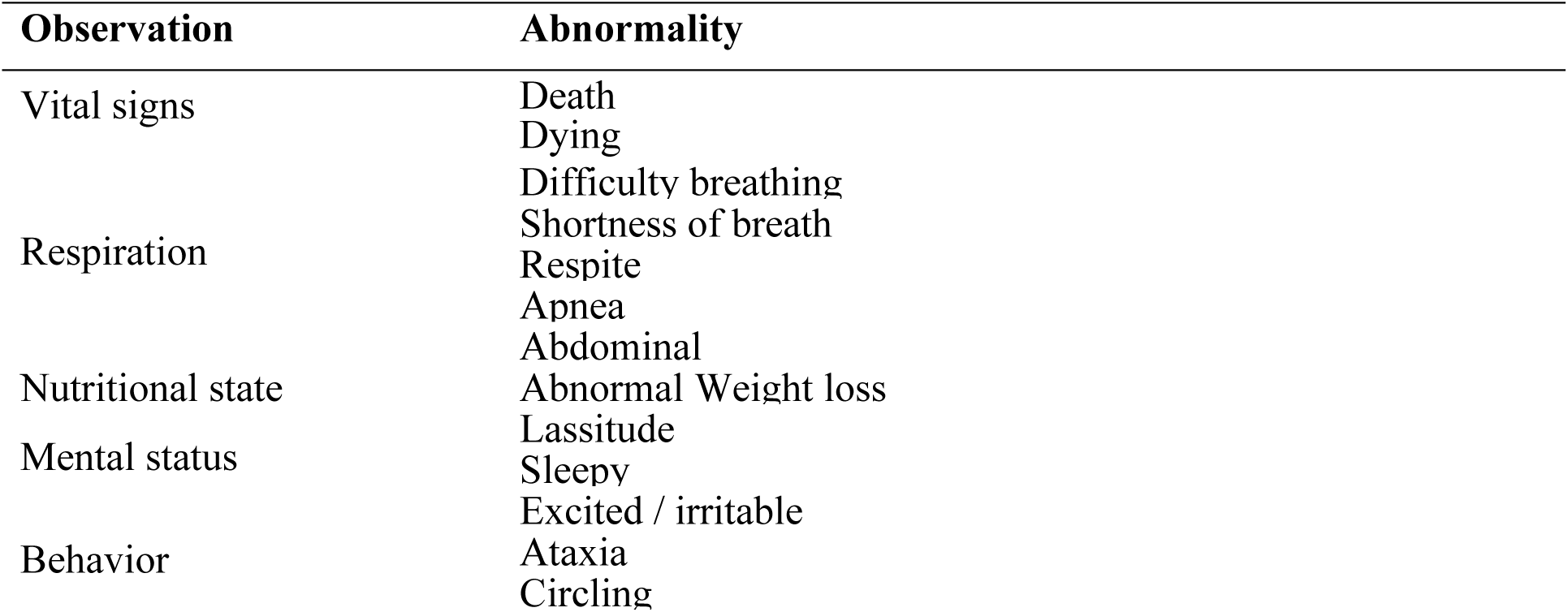
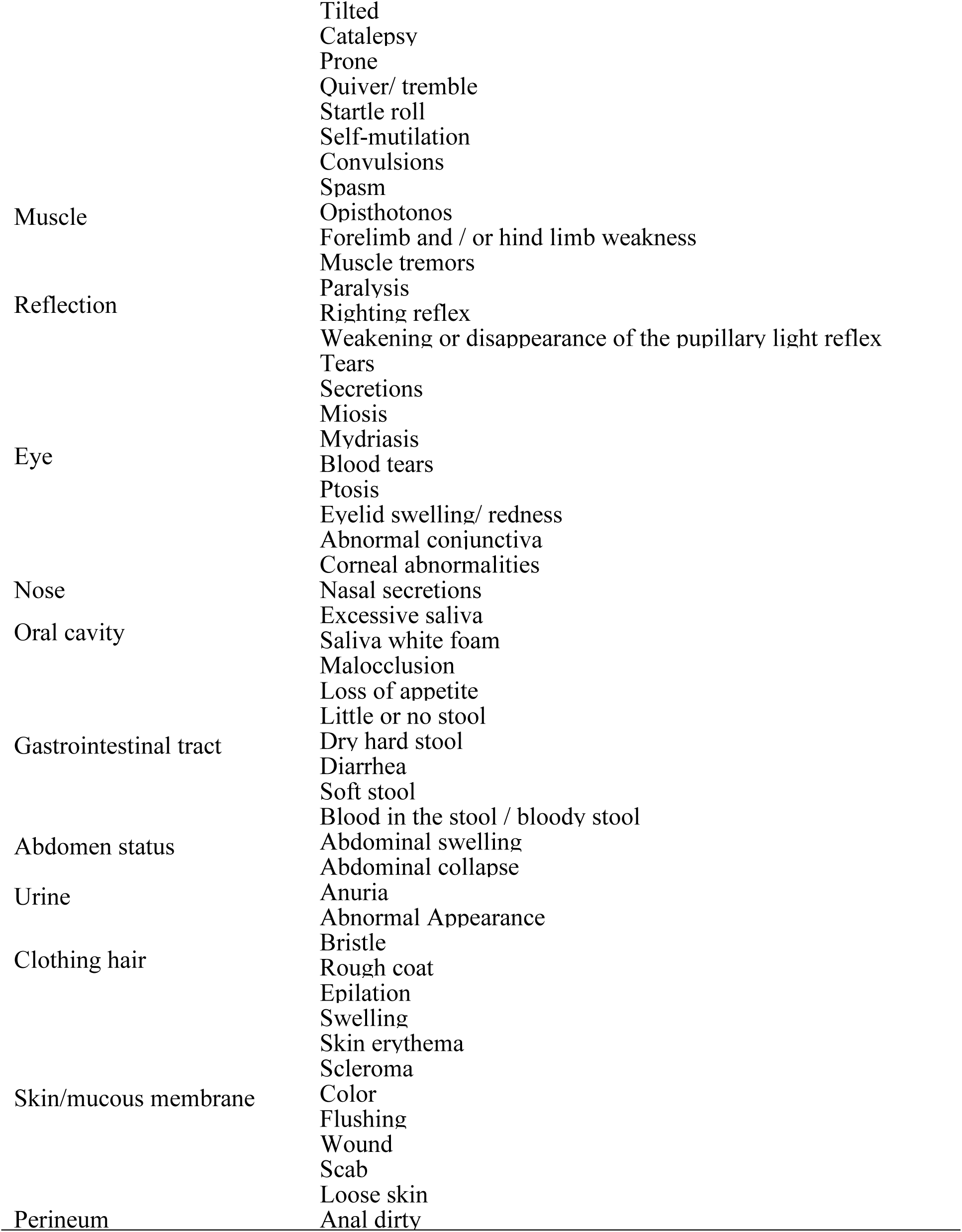
Clinical observation parameters after 4-week daily treatment in chronically dosed mice C57/BL6N. Routine observation of all animals were performed daily during the study by animal facility staff to monitor for any sick or dying animals. The detailed clinical observation items are listed below. No clinically relevant signs were noted in animals over the 4-week test article or control administrations.

**Supplementary Table 6.**
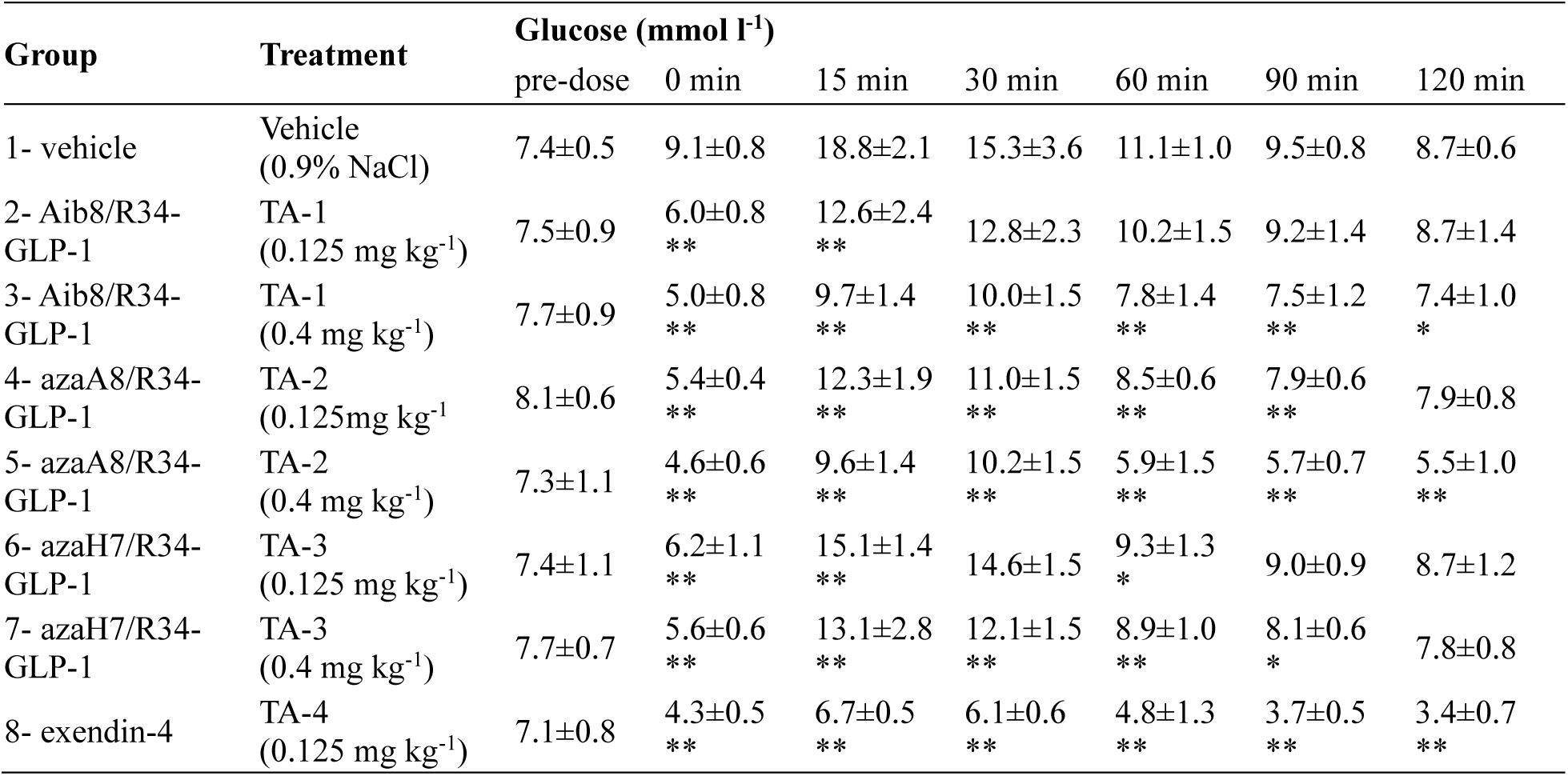
Blood glucose levels (mean ± SD) during the oGTT in 21-week-old male C57BL/6N mice after a single dose of test article or vehicle. TA-1: Aib8 (unlipidated semaglutide); TA-2: AzaA8; TA-3: AzaH7; TA-4: exendin-4. Significance vs. vehicle: *p < 0.05, **p < 0.01 (at corresponding time point).

**Supplementary Figure 1.**
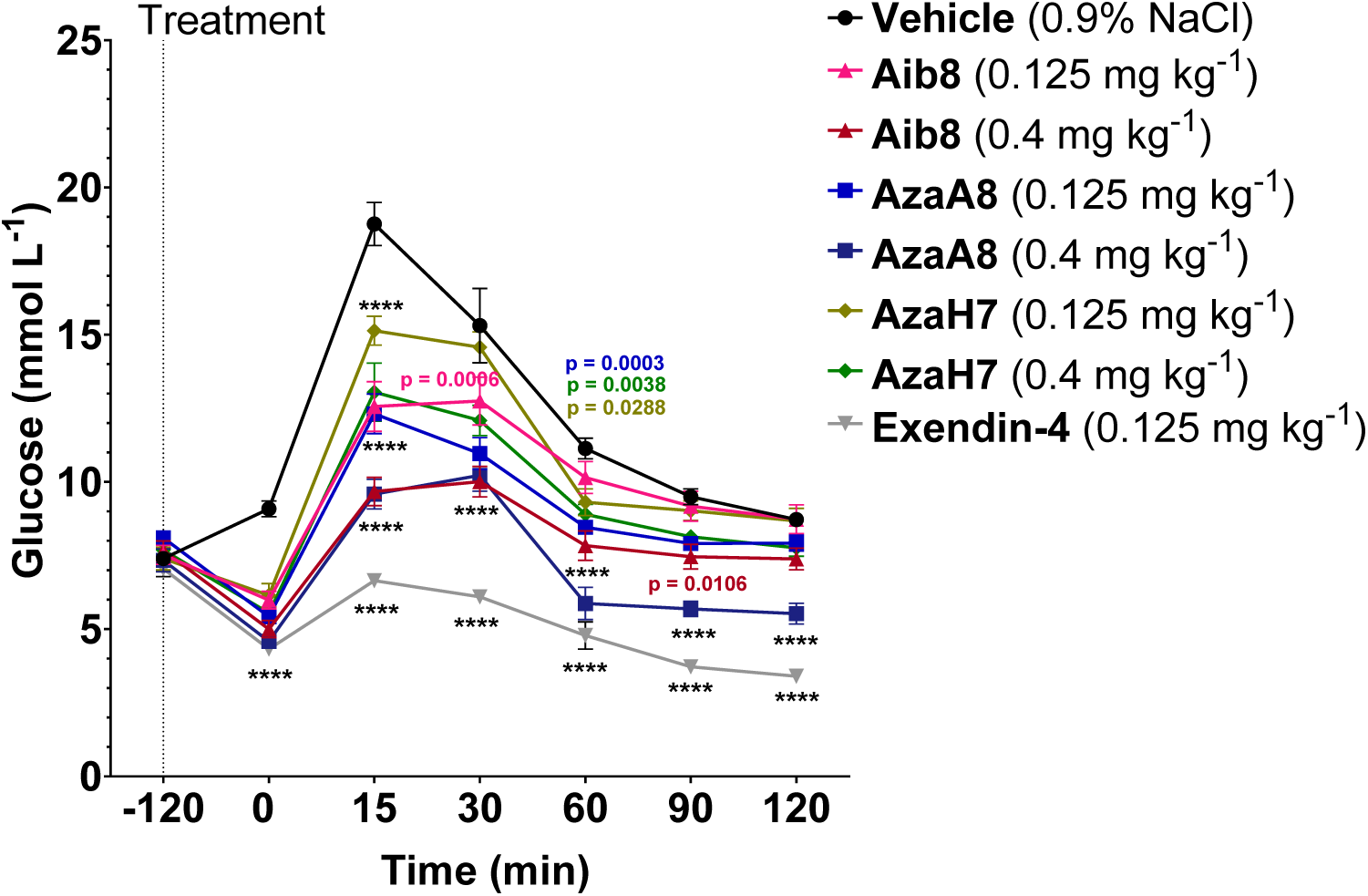
Effect of acute azapeptides treatment on glucose tolerance in lean mice (data corresponding to Fig. 3). Statistical analysis was performed using two-way ANOVA followed by Dunnett’s multiple comparisons test. P-values are indicated in the graphs and ****p < 0.0001, comparing each test article group to vehicle. When p-value is not indicated the comparison was not significant.

**Supplementary Figure 2.**
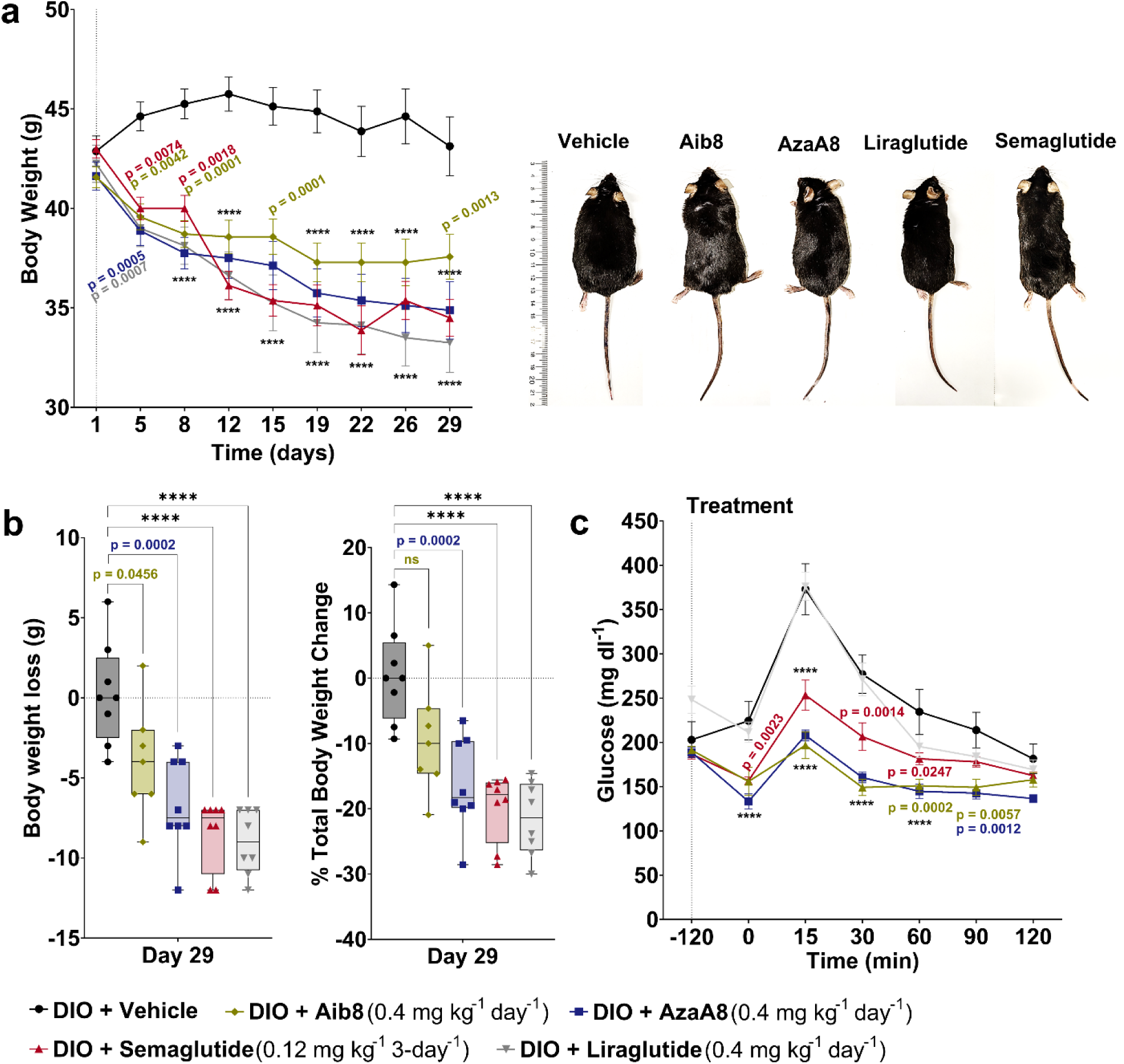
Effect of chronic intraperitoneal administration of GLP-1R agonists on body weight and glucose tolerance in DIO mice (expanded data for Figure 4). **(a)** Body weight curves (mean ± SEM) over 29 days for mice treated with AzaA8 (0.4 mg kg^-1^ day^-1^ i.p., BID), Aib8 (unlipidated semaglutide,, 0.4 mg kg^-1^ day^-1^ i.p., BID), liraglutide (0.4 mg/kg/day i.p., BID), semaglutide (0.12 mg kg^-1^ i.p. every 3 days), or vehicle (saline BID); with representative photographs of mice from the vehicle and AzaA8 groups. **(b)** Body weight loss (left), and percent (%) total body weight change (right) at endpoint (day 29) for each group. **(c)** Blood glucose levels during the oGTT at the end of treatment (day 29). *n* = 7–8 mice per group. Data are mean ± SEM. Statistical analysis: two-way ANOVA for (a) and (c) with Dunnett’s multiple comparisons vs. vehicle (ns: not significant, **p < 0.01; ***p < 0.001; ****p < 0.0001), symbols indicate the following comparisons vs. vehicle: * for AzaA8, & for Aib8, # for liraglutide, $ for semaglutide; one-way ANOVA for (b) with Tukey’s post hoc test (*p < 0.05; **p < 0.01; ***p < 0.001; ****p < 0.0001). AzaA8 and semaglutide groups showed significantly lower weights and glucose excursions compared to vehicle, whereas unlipidated semaglutide had intermediate effects. Liraglutide significantly reduced weight (similar to AzaA8) but had a less pronounced effect on oGTT glucose in this study.

**Supplementary Figure 3.**
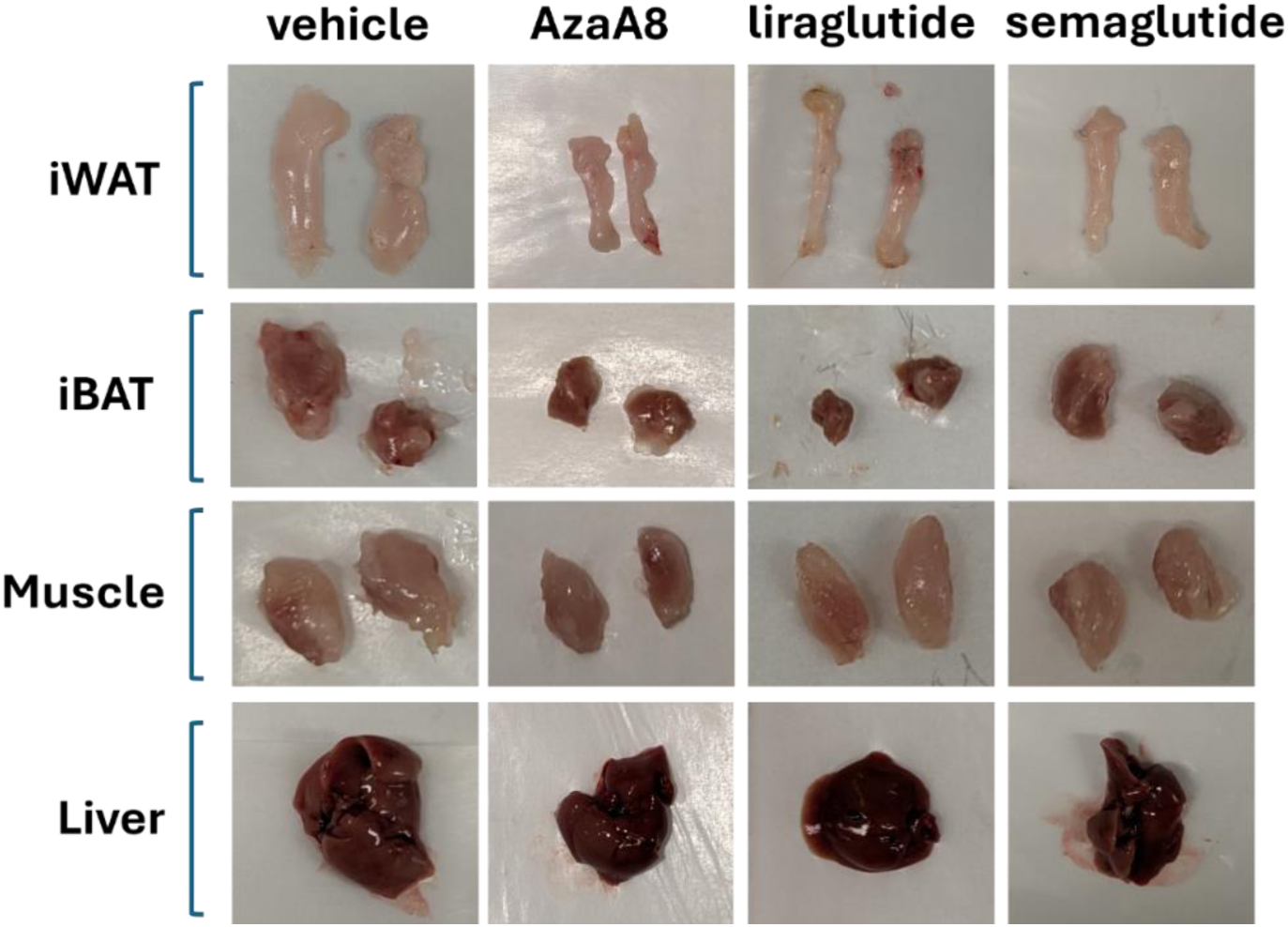
Representative tissue images from DIO mice after 29 days of i.p. treatment (corresponding to groups in. Figure 4). Shown are examples of dissected inguinal white adipose tissue (iWAT), interscapular brown adipose tissue (iBAT), quadriceps skeletal muscle, and liver from mice treated with vehicle, semaglutide, liraglutide, or AzaA8. AzaA8 and semaglutide groups exhibit visibly reduced iWAT fat pad size compared to vehicle, consistent with weight loss primarily from fat mass. iBAT and muscle tissues appear similar across groups, indicating preservation of these tissues. Liraglutide-treated mice also show reduced iWAT, but notably smaller iBAT depots than other treatment groups.

**Supplementary Table 7.** Hemoglobin A1c at endpoint of chronic treatment in DIO mice (intraperitoneal study). Values are % HbA1c (mean ±SD) measured in whole blood after 29 days of treatment. All groups maintained HbA1c in the normal range (∼4–5%), with no significant differences between treated groups and vehicle.

| **Group** | **Treatment** | **HbA1c (%) (Mean ± SD)** |
| --- | --- | --- |
| 1 | DIO + azaA<sup>8</sup>-GLP-1; weeks 1-4; 400 μg kg<sup>-1</sup> | 4.5±0.6 |
| 2 | DIO + Aib<sup>8</sup>-GLP-1; weeks 1-4; 400 μg kg<sup>-1</sup> | 4.6±0.4 |
| 3 | DIO + Liraglutide; weeks 1-4; 400 μg kg<sup>-1</sup> | 4.5±0.3 |
| 4 | DIO + Semaglutide; weeks 1-4; 120 μg kg<sup>-1</sup><br>(every 3 days) | 4.9±0.7 |
| 5 | DIO + Vehicle | 4.4±0.6 |

**Supplementary Figure 4.**
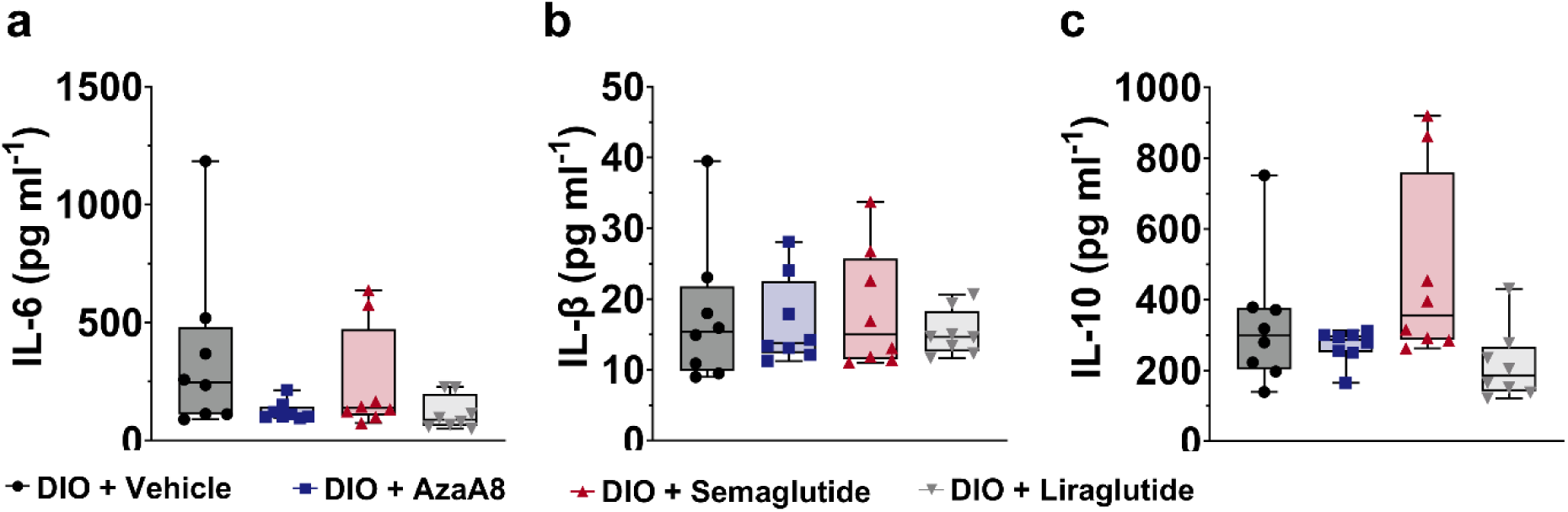
Inflammatory cytokine levels in DIO mice after chronic i.p. treatment. Serum concentrations of (a) IL-6, (b) IL-1β, and (c) IL-10 in c57BL/6N DIO mice following 29 days of treatment with vehicle, semaglutide (0.12 mg kg^-1^ Q3D i.p.), liraglutide (0.4 mg kg^-1^ day^-1^ i.p.), or AzaA8 (0.4 mg kg^-1^ day^-1^ i.p.). *n* = 7–8 per group. Data are mean ± SD. One-way ANOVA with Dunnett’s test vs. vehicle showed no significant differences (ns) for any cytokine between treatment and control groups. These results indicate that the obese mice did not exhibit elevated systemic inflammation under these conditions, and long-term AzaA8 treatment did not induce an inflammatory response.

**Supplementary Table 8.** Food consumption during chronic s.c. treatment in DIO mice. Average daily food intake (g per mouse per day) over successive intervals of the 4-week study. Data are mean ± SD. Asterisks indicate significant differences vs. vehicle over the same period (*p < 0.05; **p < 0.01 by one-way ANOVA/Tukey’s). AzaA8-treated mice showed a modest reduction in food intake during the later stages of treatment (days 15–28) relative to vehicle.

| Group | Treatment | Food Consumption(g/day) |  |  |  |  |  |  |  |
| --- | --- | --- | --- | --- | --- | --- | --- | --- | --- |
|  |  | Day 1-4 | Day 5-7 | Day 8-11 | Day 12-14 | Day 15-18 | Day 19-21 | Day 22-25 | Day 26-28 |
| 1 | DIO + AzaA8;<br>week 1 125$\mu\text{g kg}^{-1}$,<br>weeks 2-4 400 $\mu\text{g kg}^{-1}$ | 2.2±0.3 | 2.2±0.3 | 2.4±0.3 | 2.6±0.4 | 2.4±0.2<br>* | 2.3±0.2<br>** | 2.3±0.6<br>** | 2.3±0.6<br>* |
| 2 | DIO + Semaglutide<br>weeks 1-4; 40 $\mu\text{g kg}^{-1}$ | 1.0±0.4<br>** | 1.7±0.6<br>** | 2.1±0.5<br>** | 2.2±0.2<br>** | 2.1±0.2<br>** | 2.0±0.2<br>** | 2.3±0.1<br>** | 2.2±0.2<br>* |
| 3 | DIO + Vehicle | 2.5±0.3 | 2.6±0.2 | 2.7±0.5 | 2.9±0.4 | 2.8±0.4 | 2.8±0.3 | 3.0±0.3 | 2.7±0.2 |

**Supplementary Table 9.** Hemoglobin A1c percentage at terminal endpoint in s.c. chronic treatment, HFD-DIO model. C57/BL6N mice treated daily for 4 weeks were sacrificed after oGTT study and blood levels of HbA1c were measured (Pharmaron, Beijing, China) by Sjodax HbA1C test kit.

| Group | Treatment | HbA1c (%) (Mean ± SD) |
| --- | --- | --- |
| 1 | DIO + AzaA8; week 1 at 125 $\mu\text{g kg}^{-1}$,<br>weeks 2-4 at 400 $\mu\text{g kg}^{-1}$ | 3.7±0.4 |
| 2 | DIO + semaglutide; weeks 1-4 at 40 $\mu\text{g kg}^{-1}$ | 3.6±0.7 |
| 3 | DIO + Vehicle | 3.8±0.5 |

**Supplementary Figure 5.**
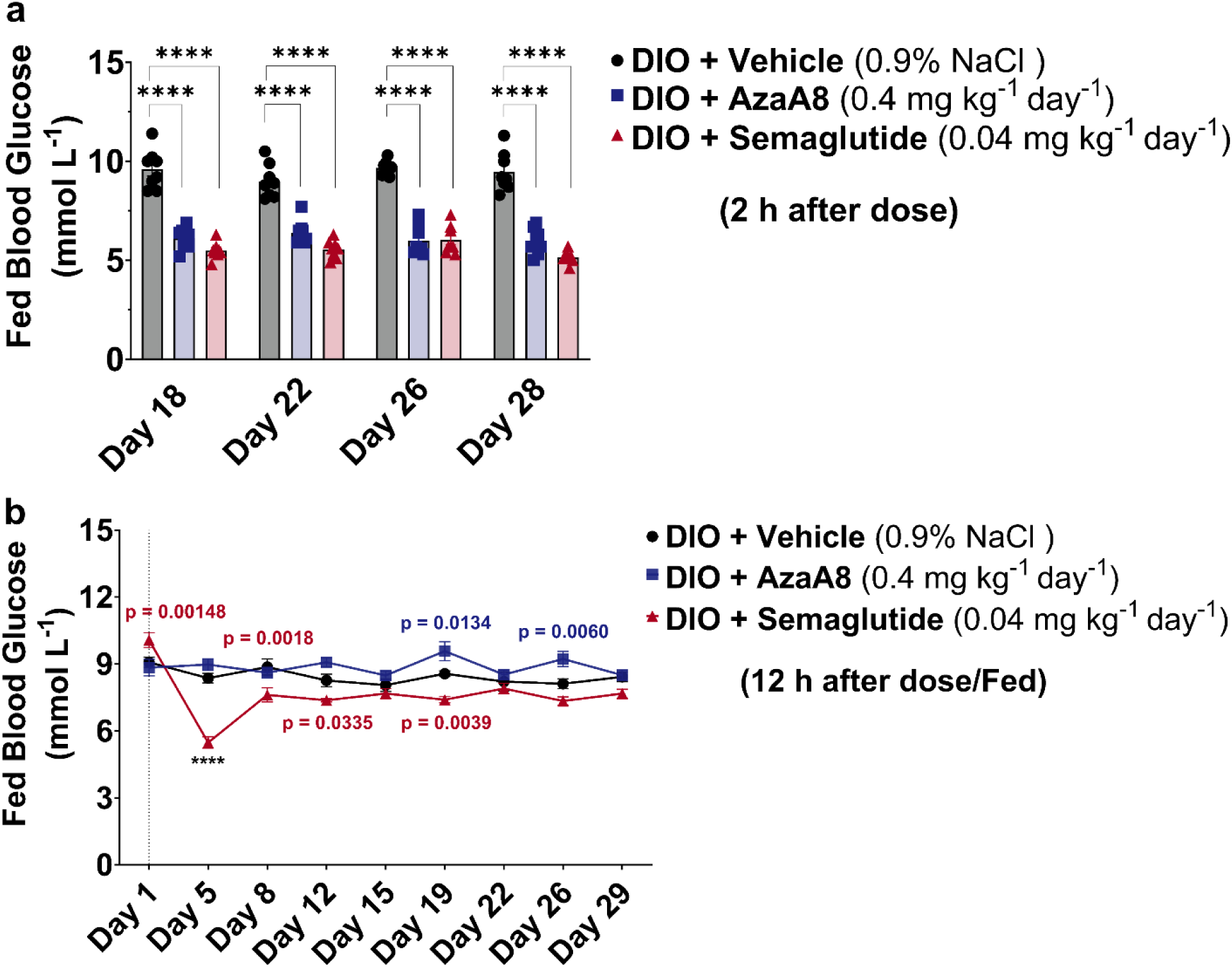
Basal (fed) blood glucose during chronic s.c. treatment in DIO mice. (a) Blood glucose measured ∼2 h after dosing (post-dose) on the indicated days during the 4-week treatment. (b) Blood glucose measured immediately before the daily dose (pre-dose baseline) on the indicated days. *n* = 8 mice per group, mean ± SEM shown. one-way ANOVA (a) and Two-way ANOVA (b) with Dunnett’s post hoc (vs. vehicle) was performed. In (a), AzaA8 (****p < 0.0001) and semaglutide (****p < 0.0001) groups showed significantly lower glucose at several time points compared to vehicle, reflecting acute on-treatment effects. In (b), no significant differences were observed between groups (all p > 0.05), indicating that baseline glycemia remained similar. Throughout the study, non-fasting glucose values in vehicle mice stayed in a relatively normal range (∼8–9 mmol l^-1^) and HbA1c values remained low (see Supplementary Table 9), consistent with the DIO model not progressing to overt diabetes in this timeframe.

